# SMAP – A Modular Superresolution Microscopy Analysis Platform for SMLM Data

**DOI:** 10.1101/2020.07.24.219188

**Authors:** Jonas Ries

## Abstract

Reconstruction of superresolved images from the raw data of blinking fluorophores in single-molecule localization microscopy (SMLM) requires extensive data analysis. Current software solutions often lack functionality, continuous development and cannot easily be extended with own algorithms. Here we release as open source a comprehensive Superresolution Microscopy Analysis Platform (SMAP) for SMLM. Its modular architecture makes it easy for anyone with basic programming experience to add own plugins, limiting development effort to implementing algorithms. It has a freely configurable graphical user interface (GUI) and currently contains more than 200 plugins spanning all steps of data analysis from single molecule fitting to post-processing, rendering and advanced quantitative analyses. SMAP is a powerful, ready-to-use software package for all steps of SMLM data analysis, that enables anyone to use state-of-the art algorithms and complex analysis workflows and that provides a versatile platform for developers to share and publish new SMLM algorithms.

---

Single-molecule localization microscopy (SMLM, such as (f)PALM^1^, (d)STORM^2^ or (DNA-)PAINT^3^) achieves superresolution imaging with nanometer resolution and can provide structural insights into cell biological questions. Reconstruction of superresolved images from the raw data of blinking fluorophores requires extensive data analysis (**Figure 1**). Free software solutions for this task exist (e.g. QuickPALM^4^, RapidSTORM^5^, Localizer^6^, ThunderSTORM^7^, Picasso^8^), but often lack continuous development and cannot easily be extended with own algorithms. Commercial software is usually bundled with a specific microscope, lacks flexibility and transparency about the underlying algorithms. This limited functionality hinders advanced quantitative analysis to extract biological insights, and has motivated many research groups to develop custom solutions (e.g. ∼100 programs participated in SMLM software challenges^9,10^). Such continuous “reinvention of the wheel” slows down progress and is a waste of limited developer resources for image analysis software, as is the need to laboriously develop various interfaces for import and export functionality and a graphical user interface with any new algorithm. Building more complex analysis pipelines requires the user to chain a variety of different programs^11^, regularly with cumbersome data conversion steps in between. Thus, all SMLM users, from newcomers to experts, would greatly profit from an open-source SMLM software platform that integrates all steps of data analysis and that can be easily used and extended.

**Figure 1:**
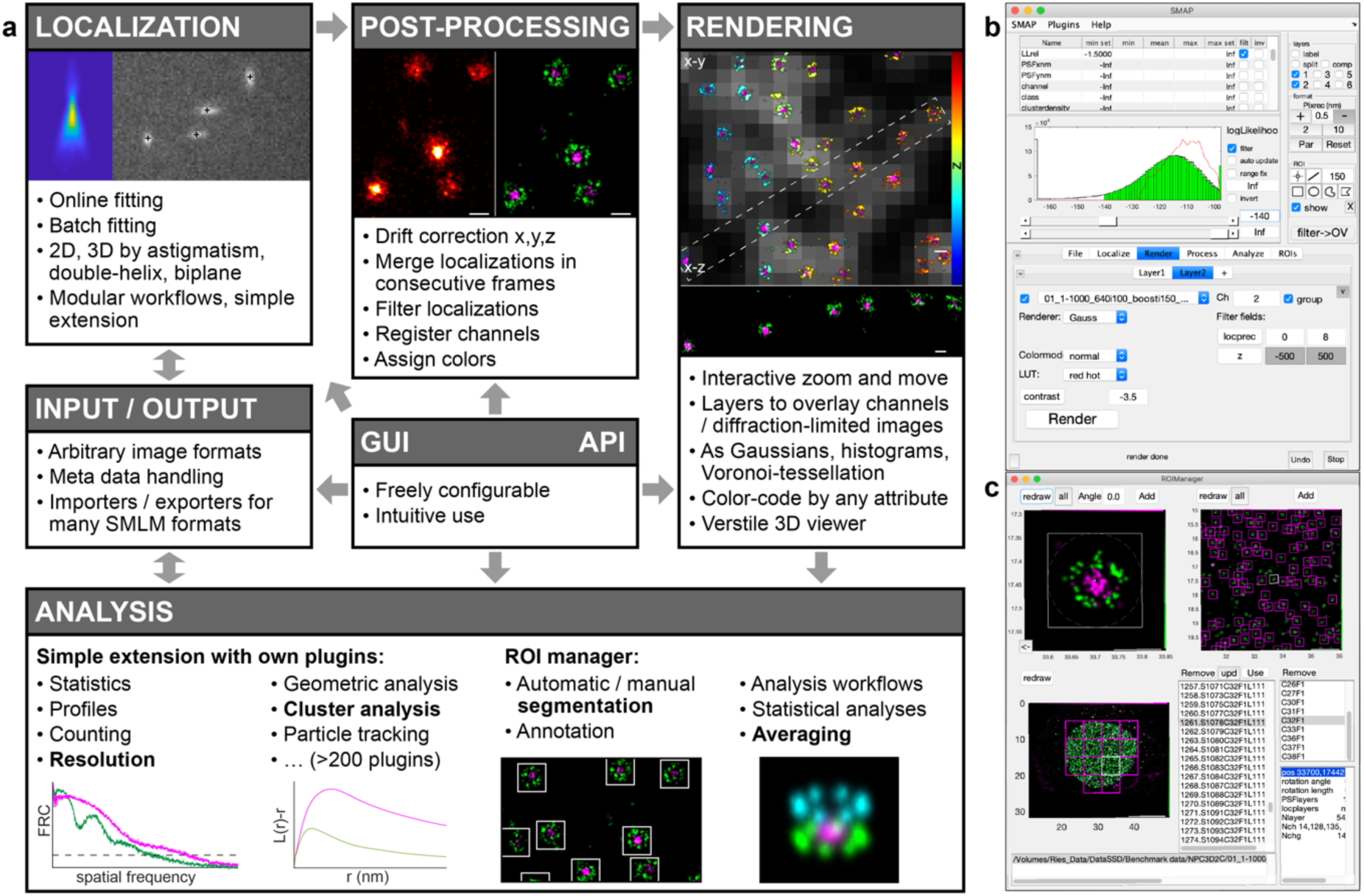
a) Overview of SMAP modular architecture and functionality. Illustrated with a dual-color 3D data set (nuclear pore complex, green: Nup107-SNAP-AF647, magenta: WGA-CF680). Error bars 100 nm. **b)** Graphical user interface of SMAP. **c)** The ROI Manager.

To address this need, we here openly release a comprehensive Superresolution Microscopy Analysis Platform (SMAP) for SMLM. Its modular architecture makes it easy for anyone with basic programming experience to add own plugins, limiting development effort to implementing algorithms. It has a freely configurable graphical user interface (GUI) and currently contains more than 200 plugins spanning all steps of data analysis from single molecule fitting to post-processing, rendering and advanced analyses (**Figure 1**). Analysis steps and parameters are logged, allowing anyone to reproduce results from raw data. In the following, we will briefly highlight the main functionalities of SMAP. A complete list of plugins can be found in **Supplementary Table 1**.

First, SMAP shows excellent performance in determining single-molecule positions^9^, which is the initial task in SMLM analysis. This can be broken down into several steps for loading of camera images, filtering, background estimation, finding of candidate molecules, fitting and saving of the results. In SMAP, each of these functionalities is encoded in a separate plugin, and many plugins are chained to workflows optimized for specific experiments. Fitting and rendering can be performed online during data acquisition, allowing users to stop their experiment when sufficient localizations have been collected or to abort imperfect measurements. A fitting workflow can be applied to many data sets for batch processing, and automatically to all new data saved in a specific location, facilitating high-throughput SMLM^12^. Individual plugins in a workflow can be easily exchanged by included or user supplied plugins, aiding development and direct comparison of specific algorithms. SMAP is compatible with most camera data formats, parses metadata and uses a configurable data base to determine camera-specific settings, greatly reducing the risk of fitting errors due to wrong parameter choices. It includes plugins for 2D and 3D fitting of unmodified or any engineered PSFs by maximum likelihood estimation using Gaussian or spline-interpolated PSF models^13^.

Second, SMAP contains various post-processing plugins to e.g. merge localizations in consecutive frames, perform 3D drift-correction, filter localizations based on their attributes or algebraic expressions thereof, register dual-channel data, or assign channels in multi-color SMLM.

Third, the in-built renderer allows for intuitive exploration of data by interactive zooming and moving. It can overlay multiple layers, each layer containing separate channels or additional diffraction limited images. The renderer supports various render modes and color lookup tables that can be also used to color-code the localizations by any attribute such as intensity, z-position, time or others. A 3D viewer is dynamically linked to a user-defined ROI and allows for real-time 3D rotations and export of animations.

Fourth, SMAP provides a large number of plugins for advanced analysis to e.g. measure resolution, plot and fit profiles, perform cluster or single-particle tracking analyses, count molecules or evaluate photophysical parameters. It contains a powerful ROI manager to automatically or manually segment structures of interest, annotate them and run them through user-defined analysis pipelines.

Such comprehensive functionality carries the risk of making a software difficult and cumbersome to use. To counteract this, SMAP includes a freely configurable GUI to give easy access to regularly used features. The GUI can toggle between an expert mode and a simple mode in which optional parameters are set to default values and hidden from sight. Expert programmers can access all functions programmatically or use the plugins directly in their own software. To facilitate extension by users with limited programming skills, SMAP is developed in MATLAB. MATLAB is among the most popular platforms for data analysis, it can be interfaced with many other programming languages, and a large fraction of algorithms for SMLM are published as MATLAB code. For users without a license we provide a fully functional stand-alone version of SMAP.

SMAP is a powerful, ready-to-use software package for all steps of SMLM data analysis, enabling anyone to use state-of-the art algorithms and complex analysis workflows. Distributed as open-source and continuously developed, SMAP additionally provides a versatile platform for developers to share and publish new SMLM algorithms.

## Supporting information

Supplementary Table: List of SMAP plugins

## Acknowledgements

We thank I. Schön, E. Klotzsch, all members of the Ries group and specifically J. Deschamps, M. Mund, U. Matti, P. Hoess Y.L. Wu, S. Liu, T. Deguchi and R. Diekmann for extensive testing of the software and contributions to the manuscript and software. This work was supported by the European Research Council (ERC CoG-724489) and the European Molecular Biology Laboratory.

## Code availability

The source code and documentation can be found at www.github.com/jries/SMAP. A compiled version for PC or Mac and example data can be downloaded at www.rieslab.de.

## References

1. Betzig, E. et al. Imaging Intracellular Fluorescent Proteins at Nanometer Resolution. Science 313, 1642–1645 (2006).

2. Rust, M. J., Bates, M. & Zhuang, X. Sub-diffraction-limit imaging by stochastic optical reconstruction microscopy (STORM). Nat. Methods 3, 793–795 (2006).

3. Jungmann, R. et al. Multiplexed 3D cellular super-resolution imaging with DNA-PAINT and Exchange-PAINT. Nat. Methods 11, 313–318 (2014).

4. Henriques, R. et al. QuickPALM: 3D real-time photoactivation nanoscopy image processing in ImageJ. Nat. Methods 7, 339–340 (2010).

5. Wolter, S. et al. rapidSTORM: accurate, fast open-source software for localization microscopy. Nat. Methods 9, 1040–1041 (2012).

6. Dedecker, P., Duwé, S., Neely, R. K. & Zhang, J. Localizer: fast, accurate, open-source, and modular software package for superresolution microscopy. J. Biomed. Opt. 17, 126008–126008 (2012).

7. Ovesný, M., Křížek, P., Borkovec, J., Švindrych, Z. & Hagen, G. M. ThunderSTORM: a comprehensive ImageJ plug-in for PALM and STORM data analysis and super-resolution imaging. Bioinformatics 30, 2389–2390 (2014).

8. Schnitzbauer, J., Strauss, M. T., Schlichthaerle, T., Schueder, F. & Jungmann, R. Super-resolution microscopy with DANN-PAINT. Nat. Protoc. 12, 1198–1228 (2017).

9. Sage, D. et al. Super-resolution fight club: assessment of 2D and 3D single-molecule localization microscopy software. Nat. Methods 16, 387–395 (2019).

10. Sage, D. et al. Quantitative evaluation of software packages for single-molecule localization microscopy. Nat. Methods 12, 717–724 (2015).

11. Sieben, C., Banterle, N., Douglass, K. M., Gönczy, P. & Manley, S. Multicolor single-particle reconstruction of protein complexes. Nat. Methods 15, 777–780 (2018).

12. Mund, M. et al. Systematic Nanoscale Analysis of Endocytosis Links Efficient Vesicle Formation to Patterned Actin Nucleation. Cell 174, 884-896.e17 (2018).

13. Li, Y. et al. Real-time 3D single-molecule localization using experimental point spread functions. Nat. Methods 15, 367–369 (2018).

